# Clinically defined mutations in *MEN1* alter its tumor-suppressive function through increased menin turnover

**DOI:** 10.1101/2022.12.20.521296

**Authors:** Suzann Duan, Sulaiman Sheriff, Uloma B. Elvis-Offiah, Travis W. Sawyer, Sinju Sundaresan, Tomasz Cierpicki, Jolanta Grembecka, Juanita L. Merchant

## Abstract

Loss of the tumor suppressor protein menin is a critical event underlying the formation of neuroendocrine tumors (NETs) in hormone-expressing tissues including gastrinomas. While aberrant expression of menin impairs its tumor suppression, few studies explore the structure– function relationship of clinical Multiple Endocrine Neoplasia, type 1 (*MEN1*) mutations in the absence of a complete loss of heterozygosity at both loci. Here, we determined whether clinical *MEN1* mutations render nuclear menin unstable and lead to its functional inactivation. We studied the structural and functional implications of three clinical *MEN1* mutations (R516fs, E235K, and A541T) recently identified in a cohort of ten patients with GEP-NETs. We evaluated the subcellular localization and half-lives of these mutated menin variants in *Men1*-null mouse embryo fibroblast cells and in hormone-expressing human gastric adenocarcinoma and murine enteroendocrine tumor cell lines. Loss of menin function was assessed by cell proliferation and gastrin gene expression assays. Lastly, we evaluated the effect of the small molecule compound MI-503 on stabilizing nuclear menin expression and function *in vitro* and in a previously reported mouse model of gastric NET development. Both the R516fs and E235K variants exhibited severe defects in total and subcellular expression of menin, and this was consistent with reduced half-lives of these mutants. Mutated menin variants exhibited loss of function in suppressing tumor cell proliferation and gastrin expression. Treatment with MI-503 rescued nuclear menin expression and attenuated hypergastrinemia and gastric hyperplasia in NET-bearing mice.

**Implication:** Clinically defined germline and somatic *MEN1* mutations confer pathogenicity by destabilizing nuclear menin expression.

## Introduction

Between 20–40% of gastroenteropancreatic neuroendocrine tumors (GEP-NETs) are associated with inactivating mutations in the Multiple Endocrine Neoplasia I (*MEN1*) gene encoding the tumor suppressor protein menin ^[1–3]^. Both sporadic and familial forms of the hereditary MEN1 syndrome mark a strongly penetrant and autosomal dominant disorder characterized by the development of tumors in the parathyroid glands, anterior pituitary, endocrine pancreas, and gastrointestinal tract ^[4–6]^. GEP-NETs that secrete the gastric peptide hormone gastrin (*i.e*. gastrinoma) comprise the most clinically aggressive of *MEN1* cases, with 60% of patients presenting with lymph node metastases upon diagnosis ^[7]^. To date, approximately 1,700 unique mutations have been identified across the coding region of the *MEN1* (11q13) locus, however mutational hotspots and a clear genotype–phenotype correlation remain ambiguous ^[8]^. Frameshift insertions and deletions comprise the majority of *MEN1* mutations (40%), while missense mutations, nonsense mutations, and splice site defects account for 25%, 10%, and 11% of the remaining mutations, respectively ^[9]^.

Originally modeled by Knudsons’ “two hit” hypothesis, complete *MEN1* inactivation leading to tumor development is thought to require an additive somatic *MEN1* mutation in the context of an existing (*i.e*. inherited) germline defect ^[10]^. However, a study of MEN1-gastrinomas showed that loss of heterozygosity (LOH) at the 11q13 locus occurred in only 46% of cases, with either partial, complete, or no LOH observed in small synchronous tumors from the same patient ^[11]^. Moreover, hyperplastic precancerous lesions in MEN1 patients with multifocal gastrin and somatostatin-secreting NETs retain both *MEN1* alleles, suggesting that an independent hit leading to allelic inactivation is a critical step in the pathogenesis of these tumors ^[11]^. These studies raise the possibility that alternative post-translational mechanisms regulate menin protein expression and function in the absence of complete LOH. Indeed, we and others have shown that menin is tightly regulated by extracellular cues including gastrin and serum-derived growth factors ^[12,13]^.

Menin negatively regulates the expression of genes involved in cell proliferation and endocrine cell specification (*e.g*., gastrin) by acting as a nuclear adaptor protein and recruiting transcriptional regulators to promoter binding sites (*e.g*. JUND and Mixed Lineage Leukemia I/MLL) ^[14–17]^. Thus, prior research has centered on mapping clinical *MEN1* mutations to specific functional domains and regulatory sequences that might preclude menin from interacting with known nuclear binding partners. Targeted deletion studies have identified two nuclear export signals (NES1 and NES2) and three nuclear localization signal sequences (NLS1, NLSa, and NLS2) that are critical for maintaining proper localization of menin to the nucleus ^[18,19]^. While aberrant subcellular expression of menin impairs its tumor suppressor function, few studies have explored the structure–function relationship of clinical *MEN1* mutations within this context.

We recently performed whole exome sequencing (WES) on a cohort of 10 patients with confirmed GEP-NETs and identified germline or somatic *MEN1* mutations in all subjects ^[20]^. We mapped these mutations to coding regions within the *MEN1* locus and focused on three mutations occurring in the central pocket and NLS regions with the potential to destabilize menin protein. Here we investigated the hypothesis that clinically defined *MEN1* mutations confer pathogenicity by destabilizing nuclear menin.

## Materials and Methods

### Plasmid Design and Purification

The coding region of the human *MEN1* gene (1830 nucleotides) was ligated to the EcoRI site of the pcDNA3.1+/C-(K)-DYK plasmid upstream of the FLAG-epitope DYKDDDDK sequence. Three single nucleotide variants previously identified by whole exome sequencing of patients with GEP-NETs ^[20]^ were introduced as follows: c.1546dupC (R516fs), c.703G>A (E235K), and c.1621G>A (A541T). As the c.1546dupC variant encodes a premature stop codon that results in protein truncation at amino acid 529, the FLAG-epitope sequence was introduced at the N-terminal domain. Plasmid construction and validation were performed by GenScript (Genscript Biotech Corp, Piscataway, NJ). BL21 (DE3) competent cells (Invitrogen) were transformed with the pcDNA empty vector or the vector expressing wild type *MEN1* containing the respective mutations. Plasmids were purified from transformed *E. coli* following ampicillin selection using the QIAGEN Plasmid Purification Midi-Prep Kit (QIAGEN, Hilden, Germany) and reconstituted in sterile Tris-EDTA (TE) buffer.

### Cell Culture

*Men1-*null mouse embryo fibroblasts (*MEF^ΔMen1^*) were a gift from Dr. Agarwal (NIH, Bethesda, Maryland). *MEF^ΔMen1^* cells were grown in Dulbecco’s Modified Eagle Media (DMEM) containing 10% Fetal Bovine Serum (FBS, Sigma Millipore), penicillin-streptomycin antibiotic. AGS and MKN-45G human gastric adenocarcinoma cell lines were purchased from ATCC (Manassas, VA). AGS cells were grown in Ham’s F12 media supplemented with 5% FBS, 5% newborn calf serum, and penicillin-streptomycin. MKN-45G cells were grown in RPMI 1640 media containing L-glutamine and supplemented with 5% FBS and penicillin-streptomycin. The previously reported on GLUTag murine enteroendocrine tumor cell line ^[21,22]^ was grown in DMEM supplemented with 10% FBS and penicillin-streptomycin. All cells were maintained at 37°C and used at passage numbers 3-10.

### Subcellular Fractionation

Within 24h of seeding cells into 6-well plates, cells were transfected with 1.2 μg of the FLAG vectors using Polyplus jetOPTIMUS reagent (Polyplus, Illkirch, France). Following 48h transfection AGS, MKN-45G, and GLUTag cell lines were washed twice in PBS and mechanically dissociated. Cells were centrifuged for 5 min at 500 x g at 4°C and then the pellet was resuspended in a hypotonic lysis buffer consisting of 20 mM HEPES, 0.1% IGEPAL, 1 mM dithiothriotol (DTT), and 1X HALT protease/phosphatase inhibitor (Thermo Fisher Scientific). The pellet was pipetted up and down 15–20X and incubated on ice for 10 min prior to pulse vortexing for 10 seconds. The cells were centrifuged for 5 min at 15,000 x g at 4°C and the suspension was collected as the cytoplasmic protein fraction. The pellet was resuspended in a hypertonic lysis buffer consisting of 20 mM HEPES, 20% glycerol, 500 mM NaCl, 1.5 mM MgCl2, 0.2 mM EDTA, 0.1% IGEPAL, 1 mM DTT, and protease/phosphatase inhibitor. The protein solution was vortexed, then incubated on ice for 1h with intervals of vortexing every 15 min. The solution was centrifuged at 15,000 x g for 15 min at 4°C and the supernatant was collected as the nuclear protein fraction. Volumes for lysis buffers were scaled at a 4:1 ratio to account for the total protein concentration in each subcellular compartment.

For MEF^*ΔMen1*^ cells, subcellular protein extracts were obtained by using the NE-PER Nuclear and Cytoplasmic Extraction Kit (Cat #78833, Thermo Fisher Scientific). Transfected cells were washed with PBS and trypsinized for 1 min using Trypsin-EDTA solution (Invitrogen). The cells were centrifuged for 5 min at 500 X g and pellets were washed with PBS prior to subcellular fractionation. Pellets were resuspended in cytoplasmic extraction reagent (CER) I buffer (0.1 mL) supplemented with protease/phosphatase inhibitor and the cytoplasmic protein fraction was obtained by following kit instructions. Nuclear extraction reagent (NER) buffer (0.05 ml) was added to remaining nuclear pellets and proteins were extracted according to manual instructions.

### Western Blot

Protein extracts (10–15 μg) were run on a precast gradient gel (4-12% Bis-Tris, Invitrogen) for either 75 min at 100 V or 65 min at 140 V. Proteins were transferred onto a PVDF membrane using the iBlot-2 transfer system (Invitrogen). The membrane was washed and blocked with 5% BSA in Tris buffered saline with 0.05% Tween-20 (TBST) buffer for 1h at 24°C. Blots were then probed overnight at 4°C with the following antibodies diluted in 5% BSA-TBST: Mouse anti-FLAG monoclonal antibody (1:10,000 dilution, Cat #F1804, Clone M2, RRID: AB_262044, Sigma Millipore), Mouse anti-Menin monoclonal antibody (1:2,000 dilution, Cat #sc-374371, Clone B-9, RRID: AB_10987907, Santa Cruz Biotechnology), Rabbit anti-Menin polyclonal (1:10,000 dilution, Cat #A300-105A, RRID: AB_2143306, Bethyl Laboratories). Membranes were washed three times in TBST, then incubated in HRP-linked anti-mouse or anti-rabbit IgG antibody for 1h at 24°C with gentle rocking (1:5,000 dilution, Cell Signaling Technology). Membranes were washed in TBST and protein bands were visualized using the Pierce ECL detection system (Cat #32106, Thermo Fisher Scientific). For loading controls, membranes were re-probed with the following antibodies diluted in 5% BSA TBST for 1h at 24°C: Rabbit anti-GAPDH monoclonal antibody (1:5,000 dilution, Cat #5174, Clone D16H11, RRID: AB_10828810, Cell Signaling Technology), Rabbit anti-β-Tubulin HRP-conjugated antibody (1:2,000 dilution, Cat #5346, Clone 9F3, RRID: AB_1950376, Cell Signaling Technology), Rabbit anti-Histone H3 HRP-conjugated antibody (1:10,000 dilution, Cat #12648, Clone D1H2, RRID: AB_2797978, Cell Signaling Technology).

For quantitation, films were scanned, converted to gray scale, and quantified using FIJI software^23^ (open source). Protein bands were identified by their expected molecular weights and quantified by equal area selection using the inverted mean gray value (MGV) method. MGVs were subtracted from 255 (pixel density maximum) to obtain the inverted MGV, then background subtraction and normalization to respective loading controls was applied. Representative images of blots were enhanced using global brightness adjustments that were applied to the entire image.

### Immunocytochemistry and Immunofluorescence Staining

AGS, MKN-45G, GLUTag, and MEF^*ΔMen1*^ cells were seeded onto glass coverslips and incubated overnight to allow the cells to adhere to coverslips. Cells were transfected with FLAG-epitope expressing plasmid using Polyplus jetOPTIMUS transfection reagent as previously described. Following 48h, cells were washed twice with PBS and fixed in 4% paraformaldehyde (PFA) for 15 min at 24°C. Cells were washed prior to permeabilization with 0.5% Triton-X in TBST for 10 min at 24°C. Coverslips were blocked for 1h at 24°C with 10% donkey serum in 0.1% BSA, 0.1% Triton-X TBST buffer, then incubated in Rabbit anti-FLAG primary antibody (1:400 dilution, Cat #14793, Clone D6W5B, RRID: AB_2572291, Cell Signaling Technology) overnight at 4°C in a humid chamber. Coverslips were washed in TBST and then incubated in Donkey anti-Rabbit IgG Alexa Fluor-conjugated 594 secondary antibody (1:500 dilution, Cat #A21207, RRID: AB_141637, Invitrogen) for 1h at 24°C in a humid chamber. Coverslips were washed in TBS, mounted with Prolong Gold anti-fade mounting medium with DAPI (Invitrogen), and imaged using the Nikon Olympus epifluorescence microscope with Cellsense software. For quantitation of nuclear and cytoplasmic expression, 10 images at 200X magnification were acquired per plasmid condition per experimental replicate. A total of 30 images per plasmid per condition were quantified and analyzed for statistical significance using Graphpad Prism 9 software (San Diego).

### Half-life Study

MEF^*ΔMen1*^ cells were seeded onto a 6-well plate at a density of 0.5 × 10^6^ cells per well. Cells were transfected 24h later with 1.2 μg of respective FLAG epitope-expressing plasmids using the Polyplus jetOPTIMUS reagent as previously described (Polyplus). After 40h, fresh media with cycloheximide was added (10 μM, Tocris). In parallel, cells were pretreated with the proteosome inhibitor MG-132 (10 μM, Cell Signaling Technology) for 1h prior to the addition of cycloheximide. Treated cells were harvested at time points ranging from 0 to 8 h. Cells were washed with ice-cold PBS and lysed in 0.1 ml of RIPA buffer supplemented with protease and phosphatase inhibitors (Thermo Fisher Scientific). Cell lysates were passed through a 27 G syringe needle 5 times, incubated on ice for 15 min, and the resulting cell homogenate was centrifuged at 10,000 X g for 15 min at 4°C. Supernatant protein samples were used for determination of the half-life by western blot as previously described.

Inverted mean grey values for protein bands were quantified as described in the previous section and expression values were normalized to respective GAPDH bands following background subtraction. The half-life was calculated by plotting normalized expression values as a function of time and fitting an exponential decay function to expression values above 0 (y = e^−kx^). The decay constant slope (k) from the fit was used to calculate the half-life by finding the time required for the signal level to drop to 0.5, expressed by the equation (x=−ln(0.5)/k), where x is the half-life ^[24]^.

### Cell Proliferation

AGS, MKN-45, and GLUTag cells were seeded at a density of 10,000 cells in 24-well plates. Cells were transfected with 0.3 μg of respective plasmid 24h post-seeding using Polyplus jetOPTIMUS reagent as previously described. Growth media was changed 24h following transfection and proliferation was assessed at 48-, 72-, 96-, and 120h post-transfection by crystal violet staining. Cells were washed with PBS prior to fixing in 4% PFA for 15 min at 24°C. Cells were washed in PBS and incubated in 0.1% crystal violet dye for 20 min at 24°C. Cells were washed with distilled water and allowed to dry prior to imaging on the ECHO Revolve inverted microscope (ECHO, San Diego CA). Crystal violet was extracted by incubating cells in 10% glacial acetic acid for 20 min at 24°C with gentle agitation. Optical density (OD) at 590 nm was measured using the Gen5 Microplate Reader and Data Analysis Software (BioTec, Dorset, UK). OD values were normalized to the empty pcDNA vector at the 48h timepoint to adjust for technical variations and cell seeding density between experimental replicates. For MEFs, the cells were transfected with respective plasmid using Polyplus jetOPTIMUS and seeded into 24-wells 24h post-transfection. Crystal violet staining was performed beginning at 48h post-seeding and up to 6 days.

### Gastrin-Luciferase Promoter Assay

The GasLuc plasmid reporter system was used to evaluate transcriptional activity at the gastrin promoter, as previously reported. A 240 base pair sequence of the human gastrin gene ^[25]^ was cloned upstream of the firefly luciferase-encoding sequence in the pGL3B reporter plasmid (Cat #E1751, Promega). Upon reaching 60–75% confluency in 6-well plates, AGS cells were co-transfected with 0.75 μg of the GasLuc plasmid construct and respective FLAG-epitope expressing menin plasmids using jetOPTIMUS transfection reagent as previously described. To account for transfection efficiency, the PRL-TK Renilla Luciferase Reporter Vector (Promega) was also co-transfected into cells. Following 24h, cells were lysed and luciferase activity was determined using the Dual-Luciferase Assay System (Cat #E1980, Promega). Luminescence was measured using the Synergy 2 Microplate Reader with Gen5 analysis software (BioTec).

### Quantitative PCR

Total mRNA was extracted from cells using the Purelink RNA Isolation kit (Invitrogen). Cells were lysed in RNA lysis buffer containing 1% beta-mercaptoethanol and homogenized by passing the lysate through a 20G syringe seven times. The lysate was mixed with equal volumes of 70% ethanol prior to adding to the Purelink spin columns for RNA extraction. Up to 1 μg of RNA was used for cDNA synthesis and quantitative polymerase chain reaction (RT-qPCR). RNA was incubated with ezDNase enzyme mix for 2 min at 37°C to digest contaminating genomic DNA prior to performing cDNA synthesis using Superscript VILO IV Master Mix (Thermo Fisher Scientific). RT-qPCR was performed using PowerUp Syber Green Master Mix with 10 ng of cDNA added per reaction (Invitrogen). RT-qPCR thermocycling conditions are as follows: 95°C for 2 min followed by 40 cycles of 95°C for 15 sec and 60°C for 1 min (BioRad). The following primers from Integrated DNA Technologies (IDT) were used at 500 nM:

*GAST*: GCAGCGACTATGTGTGTATGT (forward)
*GAST*: CCATCCATAGGCTTCTTCTTCTT (reverse)
*MEN1*: GTGCCTAGTGTGGGATGTAAG (forward)
*MEN1*: TGAAGAAGTGGGTCATGGATAAG (reverse)
*HPRT1*: TCCCAAAGTGCTGGGATTAC (forward)
*HPRT1*: CCCAGTCCATTACCCTGTTTAG (reverse)

### Animals

All animal experiments were approved by the University of Michigan’s Committee on the Use and Care of Animals. *VC:Men1^FL/FL^; Sst^−/−^*(*Men1^ΔIEC^; Sst^−/−^*) mice were generated as previously described ^[12]^. Mice were housed in a facility with access to food and water ad libitum. Experimental mice were fed omeprazole-laced chow (200 ppm, TestDiet, St. Louis, Missouri, USA) for 6 months to 1 year. Vehicle (25% DMSO, 25% PEG400, 50% PBS) or MI-503 (10, 30, and 55 mg/kg body weight) was administrated intraperitoneally, once daily for 4 weeks. Blood was drawn via submandibular bleeding prior to MI-503 administration to determine basal plasma gastrin levels and at the end of MI-503 treatment. Mouse genders used were equivalently distributed across the control and treatment groups.

### Gastrin Plasma Measurement

Mice were fasted for 16h prior to euthanasia. Blood was collected by cardiac puncture in heparin-coated tubes and plasma was collected by centrifugation at 5000 x g for 10 min at 4°C. Approximately 50 μL of the plasma was used for measuring gastrin levels using the Human/Mouse/Rat Gastrin-I Enzyme Immunoassay Kit (RayBiotech, Georgia, USA).

### Statistical Analysis

Unpaired Student’s T-test was used to test for significance between two groups. One-way ANOVA with Tukey post-test was used for comparisons of three or more groups. Two-way ANOVA with Tukey post-test was used for analysis of two or more groups when comparing more than one condition. All quantitative data reflect three or more experimental replicates, in which data were normalized to the negative control group for each experiment to account for technical variation.

## Results

### Whole exome sequencing of patient-derived GEP-NETs identifies germline and somatic mutations in the human *MEN1* gene

Whole exome sequencing (WES) was previously performed on matched buffy coat and tumor specimens from nine patients with clinically confirmed GEP-NETs ^[20]^. A tenth patient was included in the study analysis following clinical testing for the MEN1 syndrome. GEP-NETs consisted of five duodenal NETs, of which four cases were confirmed gastrinomas (DGAST), and five pancreatic NETs with one confirmed gastrinoma (PGAST). From these analyses, we identified and mapped germline mutations in nine patients and novel somatic mutations in tumor specimens collected from two patients (Figure 1). A previously reported single nucleotide polymorphism at c.1621G>A ^[26–28]^ was detected in all nine patient samples that underwent WES and occurred irrespective of tumor location. The resulting alanine to threonine missense mutation was mapped to residue 541 (A541T) immediately upstream of an accessory nuclear localization signal sequence (NLSa, aa 546–572), one of the three NLS sequences located in exon 10 (Figure We further identified frameshift mutations in two DGASTs and two PNETs spanning exons 2, 3, 8 and 10 which translated into a truncated protein. Of these, we focused on a somatic DGAST-specific single nucleotide insertion identified in exon 10 (c.1546dupC) that results in a premature stop codon at residue 516 (R516fs). In addition to these two C-terminal mutations, we included in our analyses a PNET-specific somatic missense mutation (c.703G>A) that was mapped to exon 4. This single nucleotide variant substitutes a negatively charged glutamic acid for a positively charged lysine residue (E235K) and occurs upstream of a nuclear export signal sequence (NES2, aa 258–267) (Figure 1). We next investigated whether these mutations confer altered structural and functional properties that preclude menin from functioning as a negative regulator of NET development.

**Figure 1.**
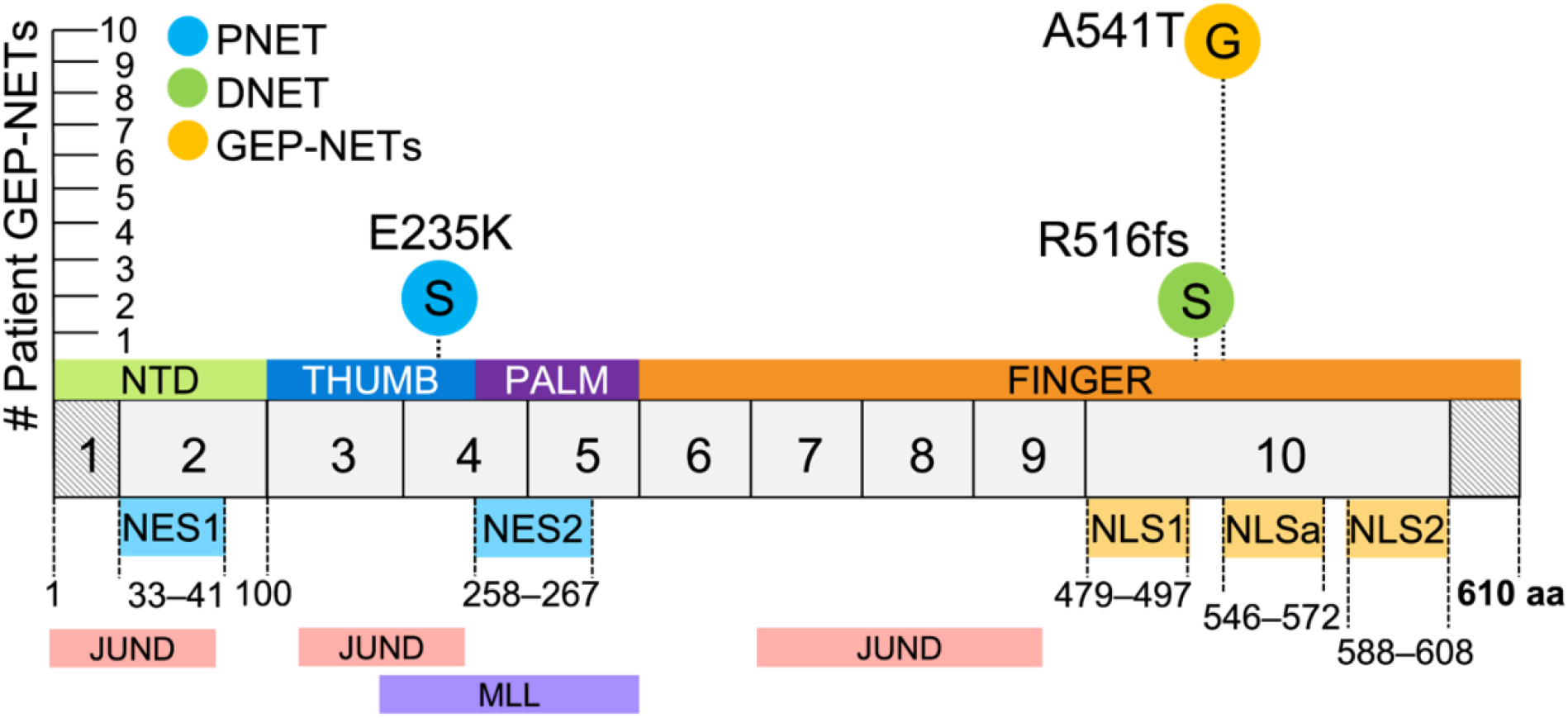
Whole exome sequencing of patient-derived GEP-NETs identifies germline and somatic mutations in the human *MEN1* gene. Germline and somatic *MEN1* mutations were previously identified by whole exome sequencing (WES) of matched buffy coat and tumor specimens from nine patients with clinically confirmed GEP-NETs ^[20]^. A tenth patient was included in the study analysis following clinical testing for *MEN1*. We focused our investigation on three single nucleotide variants: a c.1546dupC germline mutation identified in a duodenal NET (DNET) that results in a frameshift mutation and introduction of an arginine at amino acid (aa) 516 (R516fs); a c.703G>A somatic mutation identified in a pancreatic NET (PNET) resulting in a glutamic acid to lysine substitution at aa 235 (E235K); and a c.1621G>A germline polymorphism identified in all nine patient samples that underwent WES. The c.1621G>A polymorphism results in an alanine to threonine substitution at aa 541 (A541T) immediately upstream to an accessory nuclear localization signal sequence (NLSa). Menin has three NLS sequences in exon 10 and two nuclear export signal (NES) sequences. G = germline, S = somatic, aa = amino acid, NES = nuclear export signal, NLS = nuclear localization signal, NTD = N-terminal domain.

### Tumor-acquired mutations in *MEN1* result in aberrant subcellular localization at the protein level

We reasoned that clinical *MEN1* mutations occurring near the NLS and NES sequences may affect the subcellular localization of menin. As a nuclear scaffold protein, proper localization of menin to the nucleus is critical for regulating the expression of genes involved in cell proliferation and endocrine cell specification ^[18,24,29]^. To evaluate whether the R516fs, E235K, and A541T mutations altered subcellular menin expression, we constructed plasmids harboring these three mutations within the 1830 base pairs (bp) of the coding human *MEN1* sequence. The FLAG-epitope was located at the C-terminus for the two point mutations to distinguish exogenously expressed menin from the endogenous protein. Since the R516fs variant results in premature protein truncation at 529 aa, we inserted the FLAG-epitope in the N-terminal domain of the R516fs construct.

Characterization was performed in *Men1*-null mouse embryonic fibroblasts (MEF^*ΔMen1*^) to study potential structural deficits conferred by the respective mutations in the absence of any known interaction with endogenously expressed menin ^[30]^. Wild type menin and the three variants were transiently overexpressed in MEF^*ΔMen1*^ cells and protein expression was evaluated by subcellular fractionation and western blot analysis. As previously reported, wild type menin localized to both nuclear and cytoplasmic compartments, with near-equal distribution observed by biochemical fractionation (Figure 2A and 2B). A similar expression pattern was observed in cells expressing the A541T polymorphism. In contrast, the expression of both R516fs and E235K variants was uniquely restricted to the cytoplasm, with little to no expression observed in nuclear fractions (Figure 2A and 2B). Moreover, total protein abundance of these variants was significantly lower than wild type menin and the A541T mutant. Subsequent immunostaining for FLAG-epitope expression confirmed a significant increase in cytoplasmic localization of the R516fs and E235K variants compared to wild type menin and the A541T polymorphism that maintained strong nuclear expression (Figure 2C and 2D).

**Figure 2.**
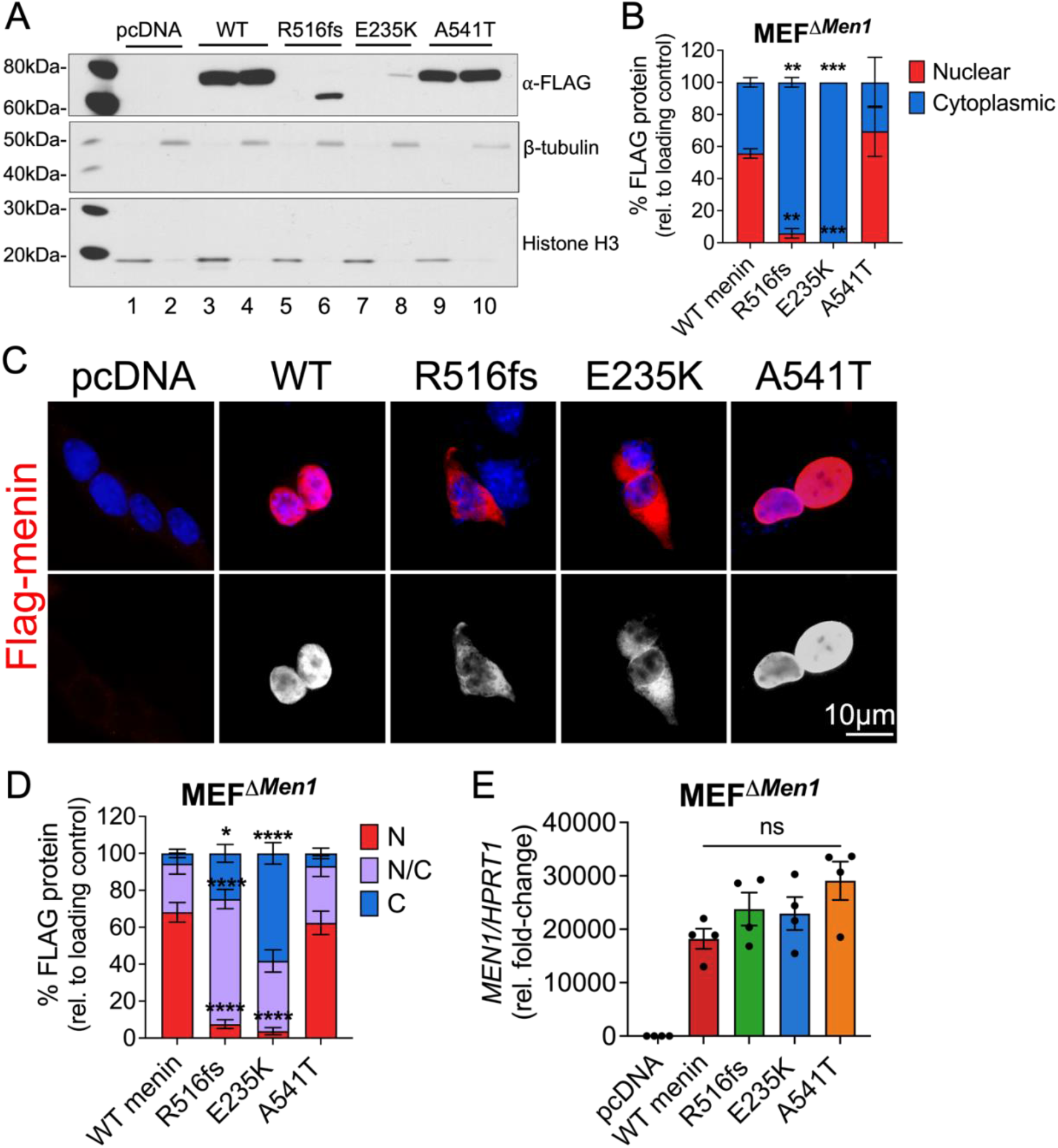
Tumor-acquired mutations in *MEN1* result in aberrant subcellular localization of menin at the protein level. Empty vector (pcDNA), wild type menin, and menin mutant proteins bearing FLAG-epitope tags were transiently overexpressed in *Men1*-null mouse embryonic fibroblasts (MEF^*ΔMen1*^). *(A)* Western blot of nuclear and cytoplasmic protein extracts probed for FLAG expression. Nuclear fractions are shown in lanes 1, 3, 5, 7, and 9 while cytoplasmic fractions are shown in lanes 2, 4, 6, 8, and 10. Histone H3 and β-tubulin were used as loading control markers for nuclear and cytoplasmic fractions, respectively. *(B)* Quantitation of FLAG protein band expression in nuclear and cytoplasmic compartments normalized to respective loading controls. n = 3 experimental replicates; ** = *p* < 0.01, *** = *p* < 0.001 by Two-way ANOVA with Tukey post-test; mean ± SEM. *(C)* Immunofluorescent images of FLAG-stained MEF^*ΔMen1*^ cells (red) co-stained with DAPI to visualize the nucleus (blue). Bottom panel shows FLAG staining in white. Scale bar = 10 μm. *(D)* Quantitation of FLAG expression in nuclear and cytoplasmic compartments expressed as a percentage of positively transfected cells. 10 images taken at 200X magnification were counted across n = 3 experiments for a total of 30 images per group; * = *p* < 0.05 by Two-way ANOVA with Tukey post-test; mean ± SEM. *(E)* Relative *Men1* mRNA expression in menin-null MEFs following overexpression of empty vector, wild type menin, and the three mutants. n = 4 experimental replicates; n.s. by One-way ANOVA.

To confirm that these effects arise from post-translational deficits as opposed to a deficiency in RNA transcription, we evaluated *Men1* mRNA levels following overexpression of wild type and mutant menin constructs. MEF^*ΔMen1*^ cells showed a ~20,000-fold induction in *Men1* mRNA expression compared to empty vector-transfected cells, and expression levels did not significantly differ between wild type menin and the mutated variants (Figure 2E). Thus, we concluded that reduced nuclear and cellular expression of the R516fs and E235K variants resulted from accelerated turnover rather than an impairment in transcription.

### R516fs and E235K mutations exhibit reduced protein stability and shortened half-lives compared to wild type menin

We next determined whether reduced and cytoplasmic expression of the R516fs and E235K variants resulted from accelerated turnover of these mutants. The half-life of menin proteins was evaluated in MEF^*ΔMen1*^ cells following transient overexpression of wild type menin or the three mutants in the presence of cycloheximide, to inhibit protein synthesis. Consistent with previous reports ^[13]^, reduced cellular expression of wild type menin was observed by 4 h (Figure 3A). Pretreatment with the proteosome inhibitor MG132 resulted in a slight reduction in the half-life of wild type menin (from 4- to 3 h), suggesting that turnover of the wild type protein was insensitive to proteosome blockade (Figure 3A and 3B). In striking contrast, expression of the R516fs variant was rapidly lost within 0.5 h of cycloheximide and pretreatment with MG132 led to a ~4-fold increase in the protein half-life (Figure 3C and 3D). Consistent with its reduced cellular expression, the E235K variant showed a significantly reduced half-life compared to wild type menin (2 h). However, unlike the R516fs mutant, proteosome inhibition did not rescue the half-life of the E235K variant (Figure 3E and 3F). Overexpression of the A541T polymorphism in the presence of cycloheximide showed this variant to be more resistant than wild type menin to protein turnover, with its half-life determined at 6 h. Moreover, pretreatment with MG132 reduced the half-life of this variant to that of wild type menin (~3.5 h) (Figure 3G and 3H). In summary, both R516fs and E235K variants exhibit a significant reduction in half-life compared to wild type menin with proteosome inhibition. There was no significant difference in the half-life of the A541T polymorphism with wild type menin. Therefore, the frameshift and exon 4 point mutations were the most unstable of the missense mutations and were potentially stabilized with proteosome inhibition. However, their incomplete response to proteosome inhibition suggested different mechanisms that mediate protein turnover.

**Figure 3.**
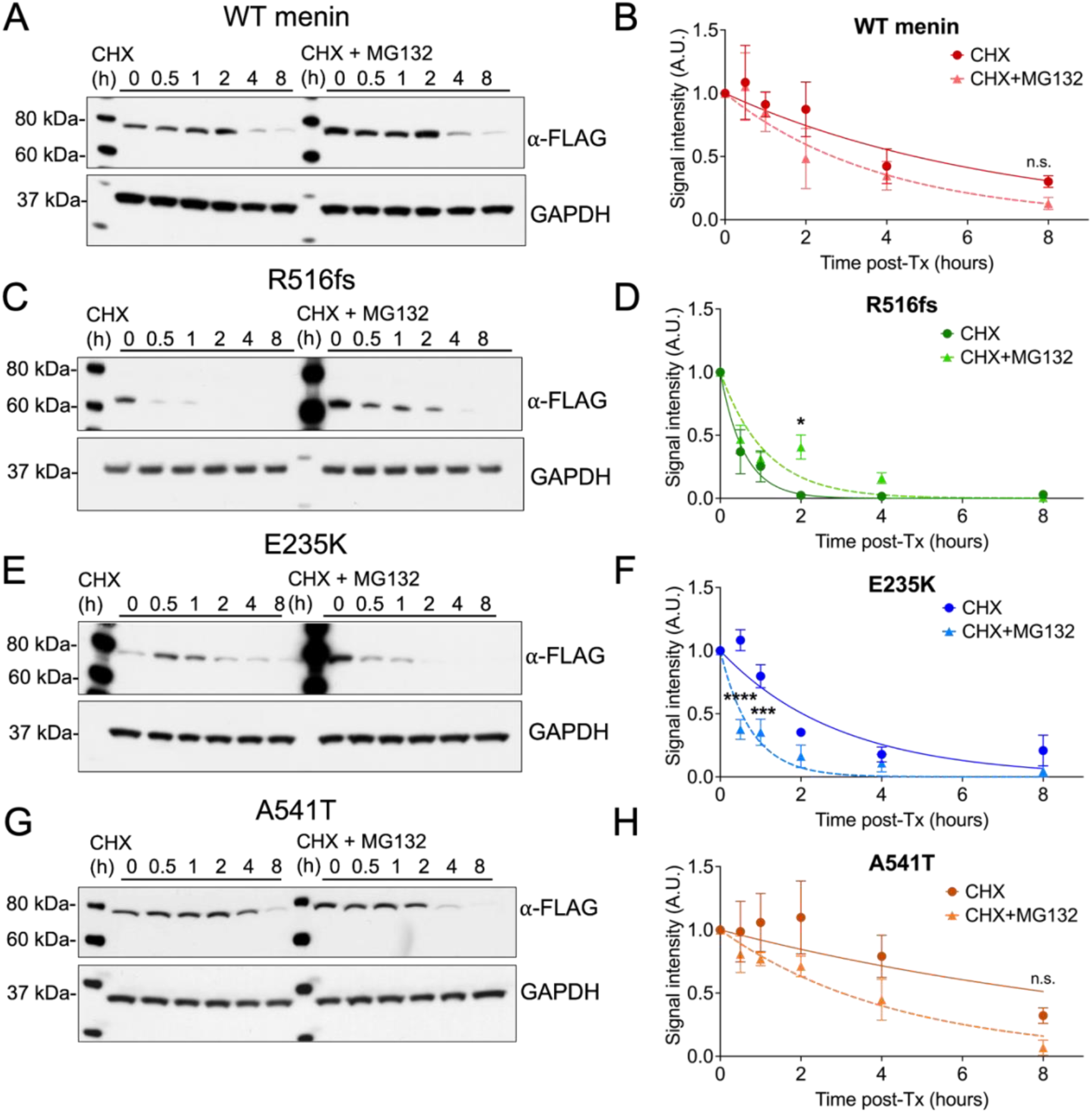
R516fs and E235K mutations exhibit reduced protein stability and shortened half-lives compared to wild type menin. *(A)* Western blot of MEF^*ΔMen1*^ whole cell extracts following overexpression of wild type menin in the presence of the protein synthesis inhibitor cycloheximide (CHX, 10 μM) alone or pretreated with the proteosome inhibitor MG132 (10 μM). *(B)* Normalized FLAG band signal intensity plotted as a function of time in the presence of CHX with and without MG132 pretreatment. *(C, D)* Overexpression of the R516fs, *(E, F)* E235K, and *(G, H)* A541T menin mutants with normalized FLAG band signal intensity plotted as a function of time in the presence of CHX alone or in combination with MG132 pretreatment. n = 3 experimental replicates, mean ± SEM. * = *p* < 0.05; *** = *p* < 0.01; **** = *p* < 0.0001 by Two-way ANOVA.

### Differential expression of menin mutants is conserved in gastrin-expressing tumor cell lines

As the previous studies examined the structural effects of *MEN1* mutations in a *Men1*-null murine stromal cell line, we next tested menin turnover in menin competent tumor cell lines. Loss of heterozygosity (LOH) at the *MEN1* locus is reported to occur in less than 50% of duodenal gastrinomas ^[11]^, thus, we performed the subsequent structure–function studies in gastric adenocarcinoma and enteroendocrine tumor cell lines expressing low levels of endogenous menin (Figure 4A). The AGS and MKN-45G human gastric adenocarcinoma cell lines were selected based on strong positive gastrin expression, whereas a gastrin-expressing subclone of the GLUTag murine enteroendocrine tumor cell line was used as model of NETs because is secretes hormones including gastrin ^[22]^.

**Figure 4.**
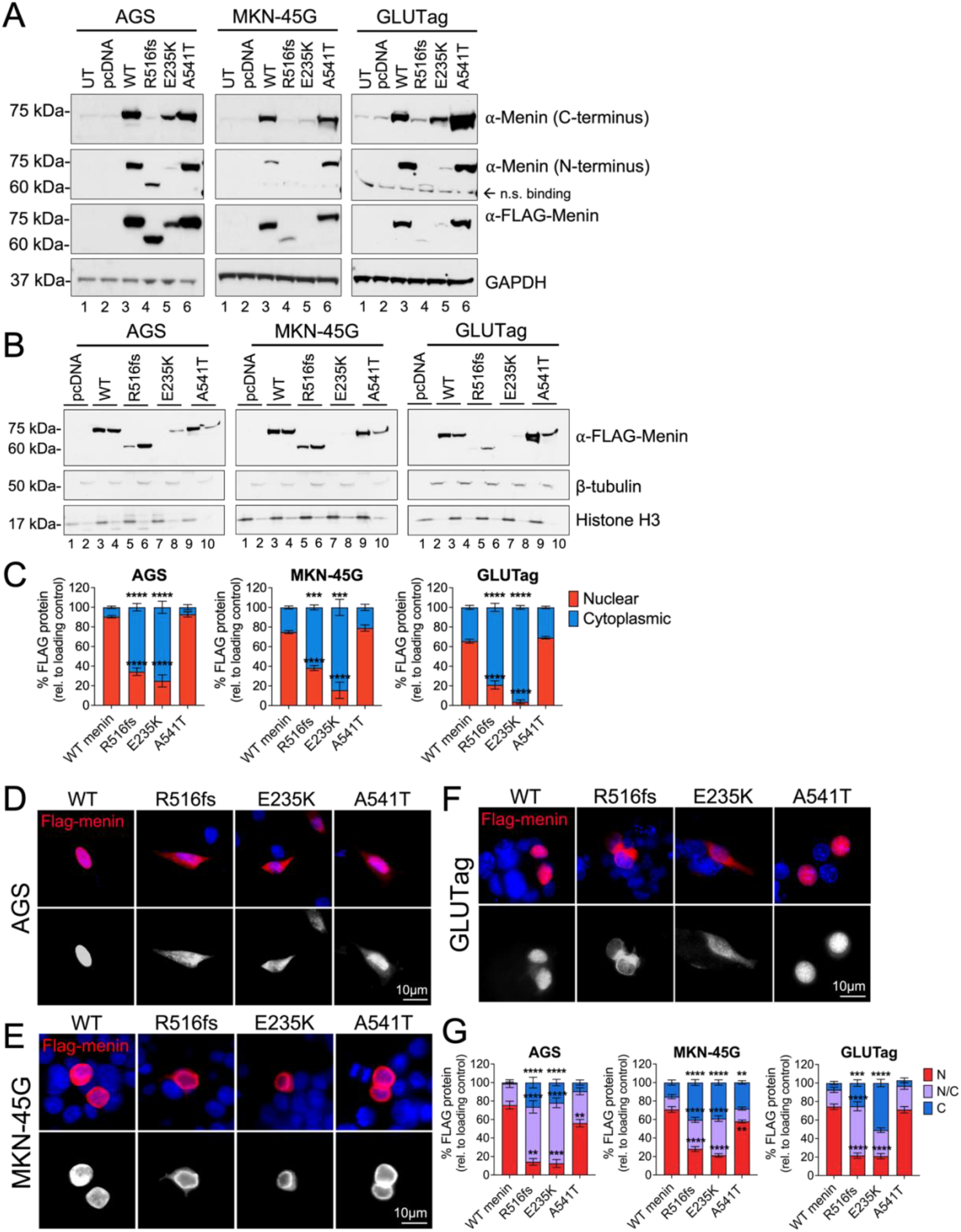
Differential expression of menin mutants is conserved in gastrin-expressing tumor cell lines. *(A)* Western blot analysis of menin and FLAG expression in whole cell extracts from AGS, MKN-45G, and GLUTag cells following overexpression of empty vector (pcDNA, lane 2), wild type menin (lane 3), and the three menin mutants (lanes 4–6). Untransfected cells (UT) were also evaluated for endogenous menin expression (lane 1). *(B)* Western blot analysis of FLAG expression in nuclear and cytoplasmic protein extracts in AGS, MKN-45G, and GLUTag cells. Nuclear fractions are shown in lanes 1, 3, 5, 7, and 9 while cytoplasmic fractions are shown in lanes 2, 4, 6, 8, and 10. Histone H3 and β-tubulin were used as loading control markers for nuclear and cytoplasmic fractions, respectively. *(C)* Quantitation of FLAG protein band intensity in nuclear and cytoplasmic compartments normalized to respective loading controls. n = 3 experimental replicates; *** = *p* < 0.001, **** = *p* < 0.0001 by Two-way ANOVA with Tukey post-test; mean ± SEM. *(D)* Immunofluorescent images of FLAG-stained AGS, *(E)* MKN-45G, and *(F)* GLUTag cells (red) co-stained with DAPI to visualize the nucleus (blue). Lower panels shows FLAG staining in white. Scale bar = 10 μm. *(G)* Quantitation of FLAG expression in nuclear and cytoplasmic compartments expressed as a percentage of positively transfected cells. 10 images taken at 200X magnification were counted across n = 3 experiments for a total of 30 images per group; * = *p* < 0.05 by Two-way ANOVA with Tukey post-test; mean ± SEM.

To further demonstrate that detection of the menin variants with FLAG and endogenous antibodies coincided, we analyzed their expression using commercially available menin antibodies raised against both C- and N-terminal peptide regions. The C-terminal menin antibody recognizes aa 575–615 ^[31–33]^, whereas the N-terminal antibody recognizes aa 1–300 of the human peptide sequence ^[31,34]^. As expected, only the N-, and not C-terminal menin antibody detected the truncated R516fs variant at a lower molecular weight compared to wild type menin and the E235K and A541T mutants (Figure 4A). Both wild type and the A541T polymorphism showed robust overexpression compared to the R516fs and E235K mutants irrespective of using menin and FLAG antibodies for protein detection (Figure 4A). Similar to the previous observations in MEF^*ΔMen1*^ cells, wild type menin protein localized to both the nuclear and cytoplasmic compartments in all three tumor cell lines, however expression was predominantly nuclear (90% in AGS, 80% in MKN-45G, 65% in GLUTag). A similar subcellular expression pattern was maintained by the A541T polymorphism. In contrast, the R516fs variant showed significantly reduced expression in nuclear fractions (35% in AGS, 40% in MKN-45G, 20% in GLUTag). Of the three mutants, the E235K variant retained the lowest nuclear expression (30% in AGS, <20% in MKN-45G, 5% in GLUTag). Immunofluorescence staining for FLAG epitope expression further confirmed predominantly cytoplasmic expression of the R516fs and E235K menin variants in AGS, MKN-45G, and GLUTag tumor cell lines (Figure 4D–G). Thus, the R516fs and E235K menin variants maintained cytoplasmic-dominant location and reduced cellular expression in tumor cell lines with low endogenous menin expression.

### Clinically defined *MEN1* mutations exhibit loss of growth-suppressive function in tumor cell lines

Following confirmation that these structural deficits were recapitulated in multiple hormone-expressing tumor cell lines, we investigated whether these clinically defined *MEN1* mutations confer pathogenic properties by inhibiting the tumor suppressive function of wild type menin. Using *in vitro* cell growth assays, we assessed whether the three menin mutants retain the ability to suppress cell proliferation. As anticipated, overexpression of wild type menin significantly inhibited the proliferation of AGS, MKN-45G, and GLUTag tumor cell lines (Figure 5A). Surprisingly, all three menin mutants showed a reduction in their ability to suppress cell proliferation, with the R516fs mutation exhibiting the most significant loss of suppressive function across AGS, MKN-45G, GLUTag, and MEF^*ΔMen1*^ cell lines (Figure 5B–E). Despite sharing similar cytoplasmic-dominant and reduced cellular expression to the R516fs variant, the E235K mutation retained a greater ability to suppress cell proliferation in all cell lines except in GLUTag cells. Intriguingly, the A541T polymorphism showed significant loss of suppressive function despite retaining similar cellular expression to wild type menin. These observations suggest a pathologic role for these clinically defined *MEN1* mutations irrespective of apparent structural deficits leading to altered subcellular expression. Moreover, these results demonstrate the potential for context-specific regulation as the apparent loss of function demonstrated by the A541T and E235K mutants was dependent on the cell type.

**Figure 5.**
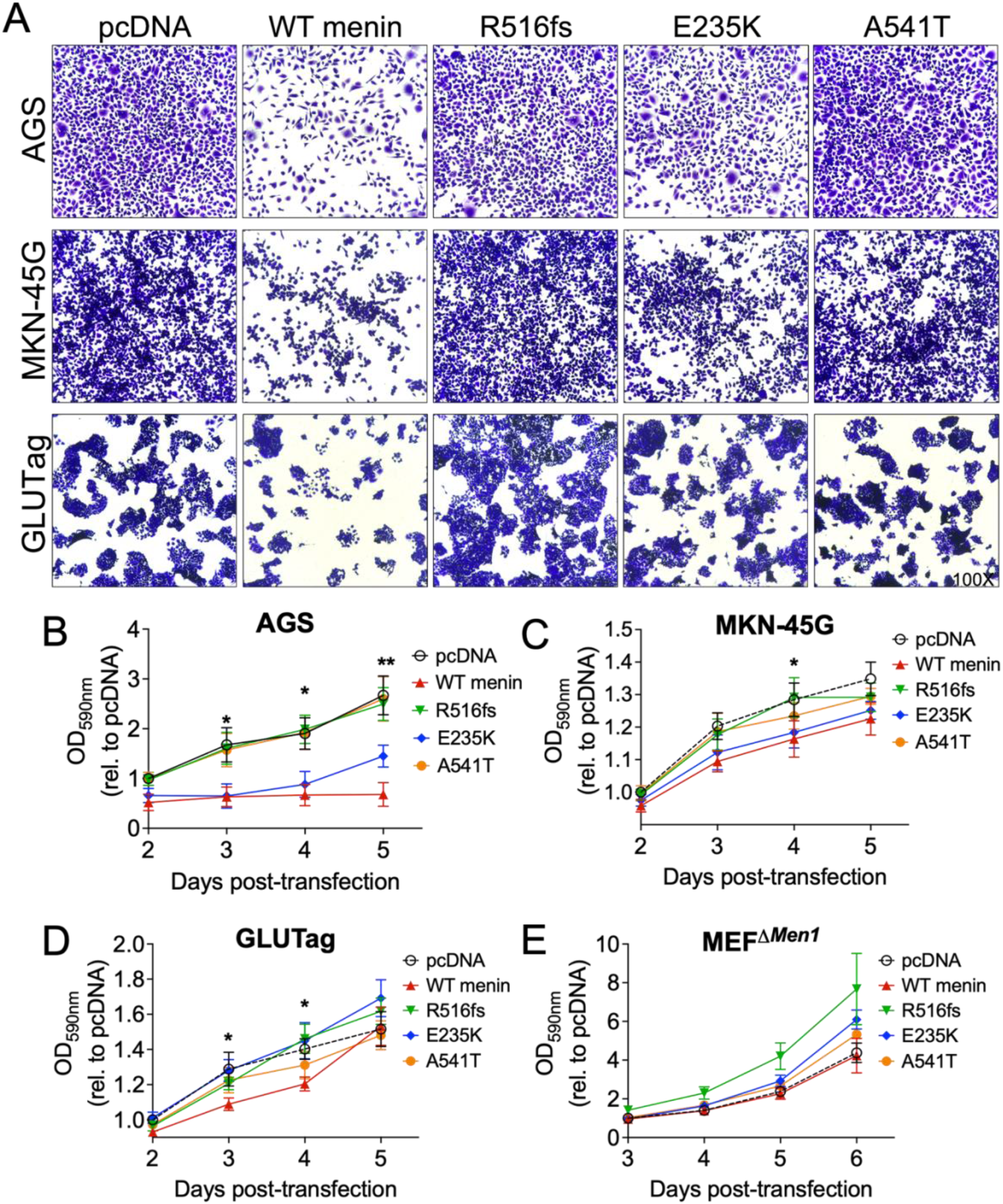
Clinical *MEN1* mutations exhibit loss of growth-suppressive function. *(A)* Representative bright-field images of crystal violet-stained AGS, MKN-45G, and GLUTag following 72 h transfection with empty vector (pcDNA), wild type menin, and the three menin mutants. Images taken at 100X magnification. (B) Normalized optical density (OD) measurements of crystal violet dye extracted from AGS, *(C)* MKN-45G, *(D)* GLUTag, and *(E)* MEF^*ΔMen1*^ cells. * = *p* < 0.05, ** = *p* < 0.01 by Two-way ANOVA with Tukey post-test; mean ± SEM.

### Clinically defined *MEN1* mutations reduce the ability of menin to repress gastrin gene expression

Consistent with its ability to repress gastrin gene expression ^[14]^, *MEN1* mutations were identified in all patients diagnosed with duodenal and pancreatic gastrinomas in our previously published cohort ^[20]^. Thus, we determined whether these clinical mutations conferred loss of function in repressing gastrin expression during the pathogenesis of these tumors. Gastrin transcript levels were evaluated in gastrin-expressing AGS and MKN-45G cell lines following transient overexpression of wild type and menin mutant proteins. As anticipated, wild type menin significantly repressed gastrin mRNA expression by 48- and 72 h in AGS and MKN-45G cells, respectively (Figure 6A and 6B). In comparison, all three menin mutations showed some degree of functional loss in repressing gastrin mRNA expression, however only the R516fs variant showed statistical significance in both cell lines (Figure 6A and 6B). We further tested the robustness of these results by assessing gastrin promoter activity using a previously characterized gastrin-luciferase (GasLuc) reporter system ^[14]^. AGS cells were transiently transfected with menin expression vectors, in addition to the GasLuc reporter plasmid comprised of 240 bp of the human gastrin gene ^[24]^ cloned upstream of the firefly luciferase (Luc) gene. Similar to the previous observations, wild type menin repressed gastrin promoter activity by 90%, however both the R516fs and A541T variants showed significant loss of function (50% repression of gastrin promoter activity) (Figure 6C). We further confirmed that *MEN1* transcript levels did not significantly differ between wild type menin and the mutated variants in AGS and MKN-45G cells (Figure 6D and 6E). Thus, loss of function was attributed to impairment in protein processing and accelerated turnover of mutated menin proteins rather than a deficiency at the transcriptional level.

**Figure 6.**
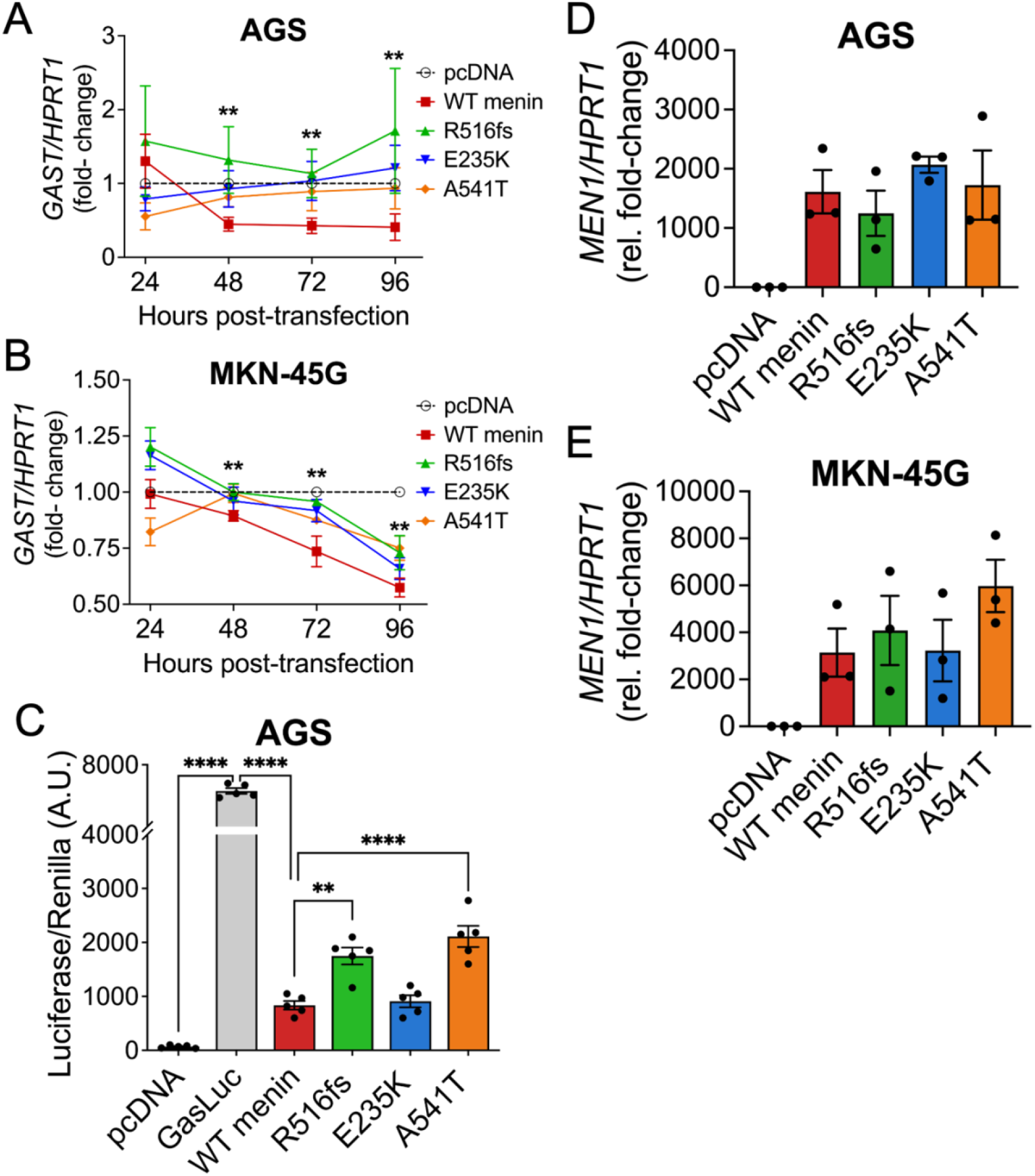
Clinical *MEN1* mutations reduce the ability of menin to repress gastrin gene expression. *(A)* Real time quantitative polymerase chain reaction (RT-qPCR) of gastrin transcript levels (*GAST*) in AGS and *(B)* MKN-45G cells following overexpression with empty vector (pcDNA), wild type menin, and the three menin mutants. *GAST* mRNA was normalized to HPRT1 expression and reported as fold-change relative to pcDNA control. ** = *p* < 0.01, by Two-way ANOVA with Tukey post-test; mean ± SEM. *(C)* The Gastrin-Luciferase (GasLuc) system was used to evaluate human gastrin promoter activity following overexpression of empty vector (pcDNA), GasLuc plasmid alone, or GasLuc plasmid in the presence of wild type menin and the three mutants. Firefly luciferase activity was normalized to the Renilla-Luciferase co-reporter. **** = *p* < 0.0001, by One-way ANOVA with Tukey post-test; mean ± SEM. *(D)* RT-qPCR of MEN1 mRNA expression in AGS and *(E)* MKN-45G cells following overexpression of the respective constructs. Not significant = *p* > 0.05, by One-way ANOVA; mean ± SEM.

### Treatment with the menin-MLL inhibitor MI-503 rescues nuclear menin expression and function

As the previously observed loss of function in the R516fs and E235K variants corresponds with impaired nuclear localization of menin, we posited whether stabilizing the expression of these mutants might rescue menin function. Menin-MLL (Mixed Lineage Leukemia, KMT2A) inhibitors (MIs), such as MI-503, are small molecule compounds that have been shown to bind directly to menin and induce thermal stabilization ^[35]^. Therefore, we tested the effect of MI-503 on menin expression in AGS cells overexpressing wild type menin and the three mutated variants. Compared to vehicle treatment, MI-503 increased the expression of nuclear menin, with the greatest effect observed in R516fs and E235K variants (~two-fold increase) (Figure 7A–C). We further evaluated whether increased menin expression rescued its ability to suppress cell proliferation. Indeed, treatment with MI-503 significantly reduced cell proliferation in the presence of R516fs, E235K, and A541T mutations (Figure 7D).

**Figure 7.**
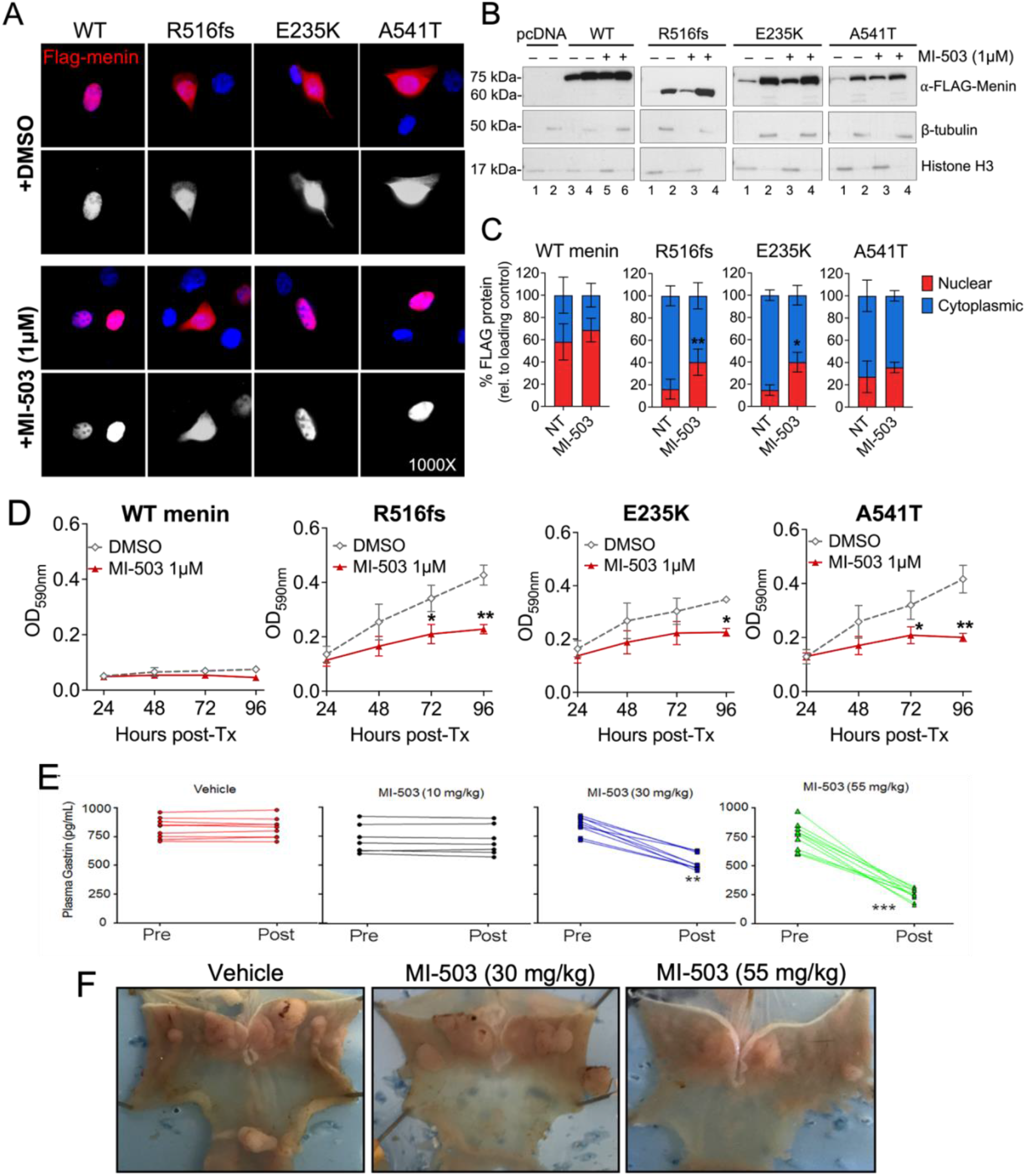
Treatment with the menin-MLL inhibitor MI-503 rescues nuclear menin expression and function. *(A)* Representative immunofluorescent images of vehicle-(DMSO) or MI-503 treated (1 μM) AGS cells expressing wild type menin and the three mutant proteins. Cells were immunostained for FLAG expression (red) and co-stained with the nuclear marker DAPI (blue). *(B)* Western blot analysis of nuclear and cytoplasmic extracts of transfected AGS cells treated with vehicle or MI-503 (1 μM). Histone H3 and β-tubulin were used as loading control markers for nuclear and cytoplasmic fractions, respectively. *(C)* Quantitation of FLAG protein band intensity in nuclear and cytoplasmic compartments normalized to respective loading controls. n = 4 experimental replicates; * = *p* < 0.05 by Two-way ANOVA with Tukey post-test; mean ± SEM. *(D)* Normalized optical density (OD) measurements of crystal violet dye extracted from transfected AGS cells treated with vehicle (DMSO) or MI-503 (1 μM). * = *p* < 0.05 by Two-way ANOVA with Tukey post-test; mean ± SEM. *(E)* Enzyme immunoassay measurement of plasma gastrin concentration from *OMS* mice before (Pre) and after (Post) treatment with vehicle, 10 mg/kg, 30 mg/kg, or 55 mg/kg of MI-503 for 4 weeks. n = 10–12 mice; ** = *p* < 0.01 by paired T-test; mean ± SEM. *(F)* Representative macroscopic images of *OMS* mice stomachs following treatment with vehicle or 30 and 55 mg/kg of MI-503. Arrows indicate to gastric NETs.

Finally, we tested whether MI-503 blocks NET development and suppresses hypergastrinemia *in vivo* by using a previously reported transgenic mouse model of gastric NET development ^[12]^. In this model, *Villin Cre; Men1^FL/FL^* mice fed an omeprazole-laced chow (OMS mice) develop hypergastrinemia and gastric NETs by 6–12 months of age ^[12]^. Three doses (10, 30, 55 mg/kg) of MI-503 were administered to OMS mice by daily intraperitoneal injections for 4 weeks. Mice exhibited a dose-dependent decrease in plasma gastrin with a maximum 70% decrease observed with the 55 mg/kg dose (Figure 7E). Consistent with reduced gastrin levels, mice showed reduced gastric hyperplasia and NET burden in the corpus (Figure 7F). Thus, MI-503 rescues the nuclear expression and function of mutated menin proteins and blocks hypergastrinemia-induced gastric NET development in mice.

## Discussion

Since its original discovery as a tumor suppressor gene ^[36, 37]^, over 1,700 mutations have been mapped to the coding region of the *MEN1* locus ^[8]^. Despite a clear association between the occurrence of inactivating *MEN1* mutations and tumor onset in cases of sporadic and hereditary MEN1 syndrome, knowledge of how these mutations manifest during pathogenesis remains limited. *MEN1* mutations are dispersed throughout its eight coding exons and fail to cluster into specific functional domains of its protein product menin, making it difficult to discern a causal relationship for the location of any given mutation. A prior study of 169 sporadic pancreatic NETs (PNETs) identified alterations in menin protein expression in the vast majority of cases (80%), with nuclear menin expression absent in tumors harboring truncating *MEN1* mutations, and strong cytoplasmic reactivity in PNETs carrying *MEN1* missense mutations ^[26]^. Given that LOH at *MEN1* loci occurs in less than 50% of MEN1-gastrinomas ^[11]^, we investigated the possibility that alternative post-translational mechanisms regulate menin protein expression in the context of incomplete LOH. We sought to determine whether clinical *MEN1* mutations predispose menin to post-translational regulation that leads to loss of nuclear expression and subsequent functional inactivation.

Here, we studied the structural and functional implications of three clinical *MEN1* mutations that we identified in a cohort of ten patients with confirmed GEP-NETs ^[20]^. Our rationale for selecting these specific mutations centered on their proximity to nuclear localization signal sequences (NLS), in addition to the presence of amino acid substitutions that confer significant structural changes or introduce the potential for post-translational modification. Of these, we examined a germline c.1546dupC frameshift insertion that leads to the expression of a premature stop codon (R516fs) before the final two NLS sequences and encodes a truncated menin protein. The c.703G>A somatic missense mutation leads to a glutamic acid to lysine exchange (E235K) and we reasoned that a substitution in polar groups might render a significant structural change near the “central binding pocket”. Lastly, the c.1621G>A germline mutation that was identified across our cohort has also been reported on extensively, with conflicting studies suggesting both a benign or a pathogenic role in predisposing patients toward NET development ^[26–28, 38]^. Approximately 1–2% of the general European population are carriers of the resulting A541T single nucleotide polymorphism (SNP), however the incidence in patients with NETs is reportedly as high as 14%, suggesting a role for this SNP in the pathogenesis of these tumors ^[26,38]^. Structurally, we posited that the exchange of an alanine to threonine residue immediately upstream to the accessory NLS (NLSa) sequence creates a potential phosphorylation site for post-translational modification. As prior studies have shown that menin is regulated by phosphorylation in response to external stimuli, such as DNA damage ^[29]^, we considered whether the A541T missense mutation might confer a similar regulatory mechanism leading to loss of nuclear menin expression and functional inactivation.

In our study, the R516fs and E235K variants exhibited severe defects in the expression and half-life of menin protein. Impaired expression coincided with a significant loss of function in suppressing cell proliferation and gastrin gene expression, with the greatest effect observed with the R516fs variant. Surprisingly, we found that the germline A541T polymorphism conferred similar loss of function despite sharing a similar expression pattern and half-life to wild type menin protein. Our findings are consistent with previous studies indicating that *MEN1* missense mutations are more susceptible to degradation compared to wild type menin protein ^[38–40]^. Moreover, we show that altered nuclear expression and loss of function exhibited by these mutants are conserved across stromal cells and hormone-expressing cell lines with epithelial and enteroendocrine characteristics. While the R516fs mutant protein demonstrated the most penetrant phenotype across all cell types examined, the E235K and A541T variants exerted differential effects depending on the cell line in which they were expressed. This raises the potential for context- and tissue-specific regulation that might inform the penetrance of inactivating mutations at different tissue sites. Indeed, our observations suggest that the E235K and A541T mutated variants may undergo degradation via a non-canonical pathway independent of ubiquitin-mediated degradation by the proteosome.

Prior studies focused on mapping structural defects at the protein level have shown that mutations arising in specific functional domains of menin can impair protein–protein interactions that may result in altered gene expression leading to cell proliferation, altered cell-specification, and NET development ^[24, 39–42]^. For instance, a recent in silico analysis of 253 disease related *MEN1* missense mutations identified three single nucleotide variants that impaired the binding affinity of menin with the Mixed Lineage Leukemia 1 (MLL) protein ^[42]^. In contrast, these mutations did not have an apparent effect on the menin–JUND interaction, further raising the potential for context-specific regulation that might dictate the tumor suppressive and oncogenic properties of menin. Given that menin is known to recruit JUND to the gastrin promoter and suppress gastrin gene expression ^[14]^, future co-immunoprecipitation studies are required to determine whether these mutations preclude menin from interacting with known binding partners, including JUND and MLL1. Finally, it should be noted that these studies are limited to standalone observations of a single given mutation. As modeled by Knudsons’ “two hit” hypotheses, examining whether germline and acquired mutations exert an additive or synergistic effect is critical to develop a comprehensive understanding of neoplastic transformation and pathogenesis.

The significance of this work is emphasized by the fact that no targeted therapeutic currently exists for patients with MEN1 syndrome. MEN1-associated neoplasms generally display a more aggressive clinical phenotype (*e.g*. multi-focal tumors), yet surveillance in this population remains insufficient as upwards of 60% of MEN1-gastrinoma cases present with metastases upon diagnosis ^[7]^. Small molecules that specifically prevent the nuclear export of menin and its subsequent degradation in the cytoplasm, in contrast to simply inhibiting the proteasome might be a promising therapeutic approach to increase nuclear menin levels and mediate suppression of gastrin expression. Menin-MLL inhibitors (MIs) are small molecules that bind directly to menin at residues F9 and P13 ^[44, 44]^ where both JUND and MLL bind. While MIs have been most extensively studied in leukemias where menin functions as a co-oncoprotein ^[45]^, MIs have also shown efficacy in blocking proliferation of solid tumors, including prostate and hepatocellular carcinoma ^[35, 46]^. Consistent with the latter studies showing MI treatment increases menin protein levels ^[45]^, we observed a similar effect in human gastric adenocarcinoma cells overexpressing mutated menin proteins. Thus, our studies support future investigation into MIs as a novel therapeutic approach for targeting GEP-NETs displaying aberrant menin protein expression in both the presence and absence of complete LOH.

## Acknowledgements

We would like to thank Dr. David Metz and Dr. Bryson Katona at the University of Pennsylvania for their assistance with acquiring the previously reported patient samples for sequencing analysis.

